# Functional diversity increases the resistance of a tritrophic food web to environmental changes

**DOI:** 10.1101/2022.07.18.500436

**Authors:** George Adje, Laurie A. Wojcik, Ursula Gaedke

## Abstract

In the light of global climate change and biodiversity loss, understanding the role of functional diversity in the response of food webs to environmental change is growing ever more important. Using a tritrophic food web model, with a variable degree of functional diversity at each trophic level, we studied the role of functional diversity on the resistance of a system against press perturbations affecting either nutrient availability or the mortality of the species, which can be interpreted as effects of eutrophication and warming respectively. We compared food webs with different levels of functional diversity by investigating the species trait and biomass dynamics, the overall changes in the species’ standing biomass as measured by the warping distance, and the duration of the system transients after the onset of a perturbation (transition time). We found that higher functional diversity increased resistance since it buffered trophic cascading effects and delayed the onset of oscillatory behaviour caused by either bottom-up forcing via perturbations to nutrient concentration or top-down forcing via perturbations to mortality rate. This increased resistance emerged from a higher top-down control of the intermediate species on the basal species. Functional diversity also promoted a higher top biomass, in particular via a higher proportion of top selective species undergoing high mortality rates. Additionally, functional diversity had context-dependent effects on warping distances, and increased transition times. Overall, this study encourages accounting for functional diversity in future investigations about the response of multitrophic systems to global change and in management strategies.

## 1 Introduction

Disturbances to ecosystems occur constantly, whether naturally or anthropogenically [Dudgeon et al., 2006, Balvanera et al., 2019], and impact their structure and functions [Brierley and Kingsford, 2009, Oliver et al., 2015], changing the surrounding environment permanently, to which the organisms must adapt. Such *press* perturbations [Bender et al., 1984, Harris et al., 2018] can have several effects on a food web. At best, the system arrives smoothly at a new static or dynamic equilibrium state which is similar to that before the perturbation [Hastings et al., 2018]. More drastically, the system may move from a static to a dynamic equilibrium, the old and new equilibria may support very different amounts of biomass at each trophic level, or extinctions could be triggered [Ives and Carpenter, 2007].

In this light, understanding the ability of a food web to cope with press perturbations is clearly of great importance. A system is considered to be *resistant* if its stable state changes little in response to a perturbation to its environment [Grimm and Wissel, 1997], considering different food web properties. A food web’s resistance to press perturbations may be assessed by considering changes to the distribution of biomass across trophic levels, to the amplitudes of biomass oscillations [Rosenzweig, 1971, Raatz et al., 2019], or to top yield and resource use efficiency (the proportion of nutrients or prey converted to top trophic level biomass) [Ceulemans et al., 2021]..

Diversity is known to influence food web stability, and consequently resistance to press perturbations, however the diversity-stability relationship is highly context-dependent [McCann, 2000, Pennekamp et al., 2018]. More recently, diversity has been considered through the lens of functional traits, which account for the role that species play within an ecosystem [Violle et al., 2007, Garnier et al., 2015]. This so-called *functional diversity* may be quantified as the difference in functional traits (such as edibility or prey selectivity) between species at the same trophic level. A few theoretical studies have demonstrated that the ability of predator and prey species to adapt their respective levels of offence and defence greatly improved resistance to perturbations [Raatz et al., 2019, Wojcik et al., 2021].

However, such studies investigating the link between functional diversity and stability have rarely considered multitrophic systems with three adaptive trophic levels [Wojcik et al., 2021], particularly in the context of press perturbations. As it is well-established that the number of trophic levels influences food web dynamics, and tritrophic sysems can be expected to show richer dynamic behaviour than simple predator-prey systems, [Matsuno and Ono, 1996, Abdala-Roberts et al., 2019], we investigated the relationship between functional diversity and resistance to press perturbations in food webs with three adaptive trophic levels.

We used a tritrophic chemostat model developed by [Ceulemans et al., 2019], in which functional diversity (modelled as a combination of prey defence and selectivity of predators to undefended prey) could be systematically varied at all trophic levels. Functional diversity influenced several ecosystem functions in this model [Ceulemans et al., 2019], as well as its resistance to a single nutrient pulse [Wojcik et al., 2021]. In this paper, we studied the effect of increasing functional diversity on the system’s resistance to environmental change, modelled as press perturbations to the nutrient inflow concentration, *N*_0_, and to the dilution rate *δ*; environmental parameters which are expected to more directly affect the bottom and the top trophic levels respectively. These changes could be caused, for instance, by an increase in temperature (a symptom of climate change), or by eutrophication (a driver of deteriorating water quality) - two urgent global issues [Brauman et al., 2019]. Increasing *N*_0_ can be thought of as eutrophication; increasing *δ* reflects, for example, warming bringing about an increased rate of turnover.

We examined the systems’ responses to press perturbations in three ways. We began by looking at the biomass and trait dynamics - that is, over which ranges a system exhibits fixed point or oscillatory dynamics, and the amplitudes of any oscillations, as well as the trophic transfer efficiency. We also quantified, via the warping distance [Raatz et al., 2019], the overall change in the attractor due to a perturbation. Finally, we considered the transition time - the time taken for a system to transition to its new attractor after the onset of a perturbation. We hypothesised that, analogously to the increase seen in resistance to a nutrient pulse [Wojcik et al., 2021], increased functional diversity would increase the system’s resistance to press perturbations.

## 2 Methods

### 2.1 Model equations

The model equations are those developed in [Ceulemans et al., 2019]: a tritrophic chemostat system with no omnivory, and with an explicitly modelled limiting nutrient pool. The system is maintained at a constant volume by removing medium (containing individuals of all species) at a dilution rate *δ* and replacing it, at the same rate, with fresh medium containing nutrients at concentration *N*_0_. The basal and intermediate trophic levels have defended (d) and undefended (u) species; the intermediate and top levels each have selectively (s) and non-selectively (n) predatory species - this means that the basal and top trophic level have two species each, the intermediate level has four (Fig. 1). Following the conventions in [Ceulemans et al., 2019], basal, intermediate, and top species biomass densities are denoted by *B, I*, and *T* respectively, and the basal species take up nutrients, with concentration *N*. Nitrogen is considered to be the limiting nutrient.

**Fig. 1.**
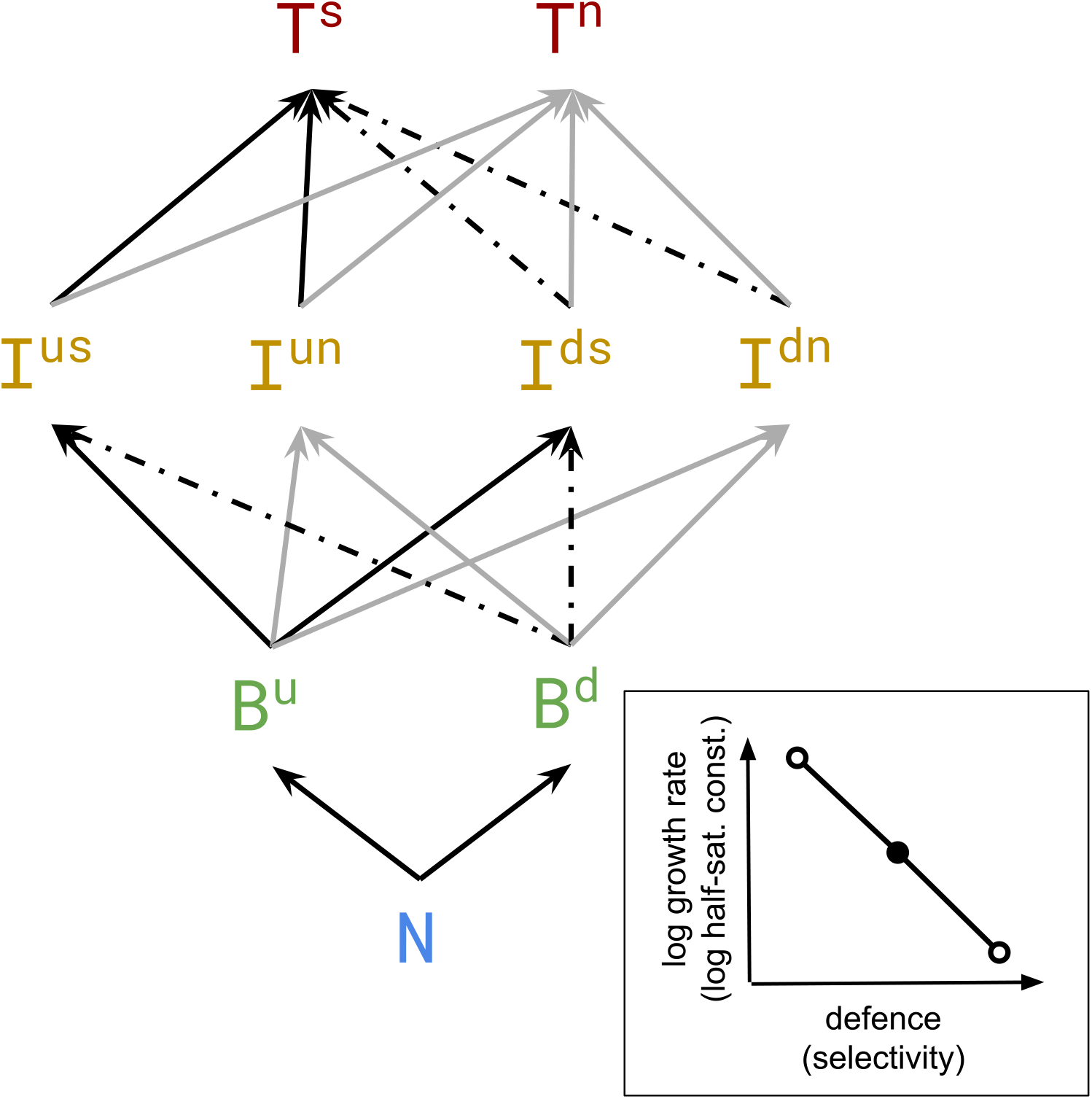
An illustration of the tritrophic food web investigated here. Nutrients *N*, basal, intermediate, and top trophic levels *B, I*, and *T* respectively. Faster-growing undefended basal and intermediate species *B*^*u*^ and *I*^*us*^, *I*^*un*^ are favoured by selective (black arrows) intermediate and top species *I*^*us*^, *I*^*ds*^ and *T* ^*s*^, which have a lower half-saturation constant than non-selectivist (grey arrows) intermediate and top species *I*^*un*^, *I*^*dn*^ and *T* ^*n*^ that are also able to feed on defended basal and intermediate species *B*^*d*^ and *I*^*ds*^, *I*^*dn*^. As Δ increases to 1, the rate at which non-selectivist species are able to feed on defended species (i.e. the interactions indicated by dotted arrows) decreases to 0, the difference between the selectivist and non-selectivist half-saturation constants increases, as does the difference between the growth rates of the defended and undefended species. Inset: an illustration of the tradeoff between growth and defence, which applies identically to selectivity and half-saturation constant. At Δ = 0, all species lie at an intermediate point *θ*_0_, indicated by the solid dot. As Δ *→* 1, the defended and undefended (selective and nonselective) species approach their extreme trait values (open circles), always remaining equidistant from *θ*_0_.

We assume that both growth/defence and selectivity/half-saturation constant tradeoffs apply: defended species have a lower growth rate, and the reduction in available resources occurring due to selective predation is offset by the selective predators’ having a lower half-saturation constant (Fig. 1).

The equations of the model are as follows:

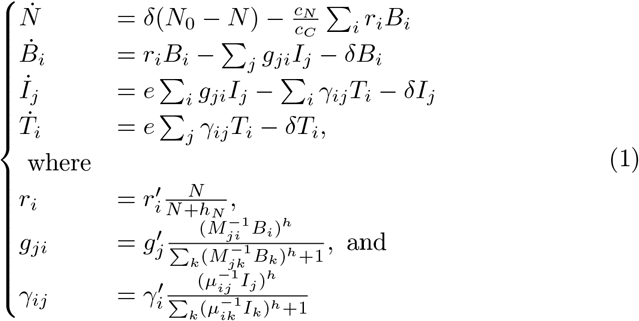

As biomass is measured in carbon-based units, the nitrogen-to-carbon weight ratio 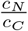 is used to scale the second term in the expression for 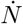. The basal growth rates *r*_*i*_ are described by a Monod function with nutrient half-saturation *h*_*N*_ ; the intermediate and top grazing rates *g*_*ji*_ and *γ*_*ij*_ are described by Holling type III functional responses, as is suitable for plankton food webs [Uszko et al., 2015], both with Hill coefficient *h*.

The ratio of the body masses between trophic levels is denoted by *m*, and the growth rates (and therefore the grazing rates) of the intermediate and top trophic levels are scaled allometrically with body mass, with scaling exponent *λ*. The remainder of the freely-adjustable symbols used in the expressions above are defined in Table 1.

**Table 1.** Parameters, symbols, and values used for each parameter in the tritrophic food web model under consideration. The values are based on, but differ slightly from, those given in [Ceulemans et al., 2019]. ^*^For perturbations to *δ, N*_0_ was set to 1200*µg/l*. ^†^For perturbations to *N*_0_, *δ* was set to 0.055 l/day.

| Parameter | Symbol | Value |
| --- | --- | --- |
| undefended basal max. growth rate | $r'_u$ | 1 /day |
| defended basal max. growth rate | $r'_d$ | 0.67 /day |
| intertrophic body mass ratio | m | 100 |
| nitrogen-to-carbon conversion ratio | $\frac{c_N}{c_C}$ | 0.175 |
| allometric scaling exponent | $\lambda$ | -0.15 |
| nutrient half-saturation constant | $h_N$ | 10 $\mu gN/l$ |
| nutrient inflow concentration | $N_0$ | 150 - 2000* $\mu gN/l$ |
| intermediate non-selectivist half-saturation constants | $M_g$ | 300 $\mu gC$ |
| intermediate selectivist half-saturation constants | $M_s$ | 150 $\mu gC$ |
| top non-selectivist half-saturation constant | $\mu_g$ | 400 $\mu gC$ |
| top selectivist half-saturation constant | $\mu_s$ | 200 $\mu gC$ |
| conversion efficiencies | e | 0.33 |
| dilution rate | $\delta$ | 0.01-0.11<sup>†</sup> 1/day |
| Hill coefficients | h | 1.2 |

The functional diversity present in the model is controlled by the parameter Δ, which varies between 0 and 1. As Δ is varied, the logarithms of the maximum growth rate, defence level, and half-saturation constant, selectivity level vary linearly: if *θ* is any of these parameters, then

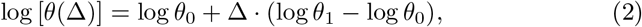

where *θ*_1_ and *θ*_0_ are the values of *θ* when Δ = 1 and Δ = 0 respectively. Fig. 1

Where Δ = 1, the food web is at its most functionally diverse: defended prey are consumed exclusively by non-selective predators, and the differences between the respective growth rates and levels of defence of the prey species are at a maximum, as is the difference between the half-saturation constants and the level of selectivity of the predatory species. In contrast, setting Δ = 0 describes the situation in which all species on a trophic level are functionally identical: in this case, the log of each trait parameter is the average of the logs of the two extreme values of the parameter in the food web in which Δ = 1 (see the comparison between the unfilled dots representing the maximally different species at Δ = 1 and the filled dot at Δ = 0 along the linear tradeoff in Fig. 1). Concretely, using growth rate as an example, this means:

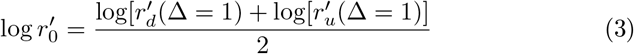

where 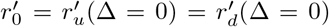 is the value of *r*′ taken on by all basal species at Δ = 0. Therefore, Δ = 0 corresponds to a simple chain, while Δ = 1 corresponds to a maximally diverse food web. For a detailed description of the model, please see [Ceulemans et al., 2019].

### 2.2 Press perturbations

For a range of values of Δ, an initial time series was obtained from the model, starting from a point *x*_0_ found, by brute force, to lie in the basin of attraction of a monostable static or dynamic equilibrium. After removing the transient dynamics, a number of points *x*_1_ on the resulting pre-perturbation attractor were randomly selected, and used as starting points for second simulations with a modified perturbation parameter. A modification of the model parameters in this way corresponds to a so-called press perturbation - an instantaneous, permanent change in environmental conditions. Regions of ecologically reasonable parameter space were selected such that the perturbation parameter could be varied across a sufficiently large range while constraining the dynamics of the system to fixed points and limit cycles, and avoiding extinctions and chaos.

Two environmental parameters were separately perturbed: the nutrient inflow concentration *N*_0_, and the dilution rate *δ*. These were chosen as two variables which affect the system most readily from opposite ends: changes to *N*_0_ directly affect the basal level, whereas changes to *δ* have the strongest immediate effect on the top level due to the top species’ having the slowest growth rates. The system can therefore be subjected to eutrophication by an increase to *N*_0_. On the other hand, changing the dilution rate *δ* is to change the rate of turnover of the entire system in the context of an increase in temperature, for instance.

The solutions of the differential equations were obtained numerically in C using the SUNDIALS CVODE solver [Hindmarsh et al., 2005], with relative and absolute tolerances of 10^−10^. Output data were studied using Python and its scientific computing libraries NumPy, SciPy, and Matplotlib [Harris et al., 2020, Virtanen et al., 2020, Hunter, 2007].

### 2.3 Measures of resistance and trophic transfer efficiency

Due to the fact that press perturbations were being applied to the system the resistance of the system was a natural measure to investigate: that is, the degree to which the perturbation causes the system to change. Since top yield is often a quantity of interest from the point of view of ecosystem functions, the efficiency of trophic transfer to the top trophic level was also considered. In the following paragraphs we describe the techniques employed to quantify both resistance and trophic transfer efficiency.

#### 2.3.1 Trophic level biomasses

The temporal mean, median and quartiles of biomass at each trophic level were compared for different values of Δ and at different values of the perturbation parameters to investigate the impact of functional diversity.

To compare systems with different levels of functional diversity, time series were produced in nine dimensions; that is nutrient, two basal species, four intermediate species and two top species. Note that when Δ = 0, species within each trophic level have identical traits, and the system is equivalent to a food chain. Therefore, the model is effectively four-dimensional, corresponding to three trophic levels and the nutrient pool. The comparison between a food chain and food webs with higher levels of functional diversity is nonetheless meaningful: when the trait values of all species at the same trophic level are identical, the system’s equations can be rewritten by introducing new state variables *B, I*, and *T* such that *B* = *B*^*u*^ + *B*^*d*^, *I* = *I*^*us*^ + *I*^*un*^ + *I*^*ds*^ + *I*^*dn*^, and *T* = *T*^*s*^ + *T*^*n*^, which allow for the recovery of the equations for a four-dimensional food chain, as detailed in previous studies using the same model [Ceulemans et al., 2019, Wojcik et al., 2021] and justified by [Moisset de Espanés et al., 2021]. In addition, all calculations were performed at the trophic level (rather than the species level) to unify the comparison.

To shed light on the underlying reasons for the results obtained, the same summary statistics were calculated for trait proportions, i.e. the ratios of defended to undefended basal and intermediate species, and of selective to non-selective intermediate and top species, at each (static and dynamic) equilibrium reached over a range of *N*_0_ and *δ*.

According to classical chemostat theory [Smith and Waltman, 1995, Harmand et al., 2017], increasing the nutrient inflow concentration (dilution rate) will lead to a corresponding increase (decrease) in top predator abundance. However, depending on Δ, a given perturbation may alter the top trophic biomass differently. Considering the proportion of basal biomass taken up by the intermediate level (as opposed to washed out), and in turn, the proportion of intermediate biomass taken up by the top level allowed for a quantification of the efficiency of biomass transfer to higher trophic levels. If B, I, and T are the total basal, intermediate, and top trophic level biomasses, *B*_*up*_ and *I*_*up*_ are the amounts of basal and intermediate biomass respectively reaching the next trophic level, *B*_*out*_, *I*_*out*_ and *T*_*out*_ are the amounts of the different trophic level washed out from the system, and *B*_*eff*_ and *I*_*eff*_ are the biomass transfer efficiencies of the basal and intermediate levels, then:

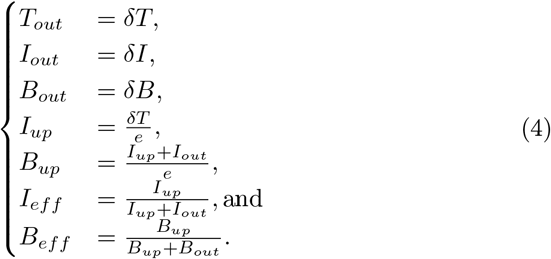

#### 2.3.2 Warping distance

In order to quantify the effect of a press perturbation on the dynamics of the food web as a whole, the *warping distance* between the pre- and post-perturbation attractors, as outlined in [Raatz et al., 2019] was calculated. The warping distance quantifies how a pair of periodic attractors differ in their positions in phase space, and in their oscillation amplitude.

Specifically, the warping distance between two periodic attractors A and B is obtained by first identifying a section of each time series corresponding to a single period of each attractor. Between two fixed points, the warping distance reduces to the distance between the two points. Following [Raatz et al., 2019], the period of a single component of the time series was taken to be the mean of the time between like peaks, which were found using `scipy.signal`’s `find_peaks` function with the `height` parameter set to 90% of the maximum peak height of the time series. The overall period of the attractor was then taken to be the lowest common multiple of the periods of oscillation of the nine individual components of the food web.

Species biomasses were summed at the trophic level to create a four dimensional system, so that each point on the attractor could be represented as a point in 4-dimensional log-biomass (equivalent) phase space. For each point *p*_*A*_ on the pre-perturbation attractor A, the point *p*_*B*_ which is its nearest neighbour on the post-perturbation attractor B was ascertained. Similarly, for each point *q*_*B*_ on attractor B, the point *q*_*A*_ which is its nearest neighbour on attractor A was ascertained. Figure 7 in the Appendix illustrates this procedure.

**Fig. 2.**
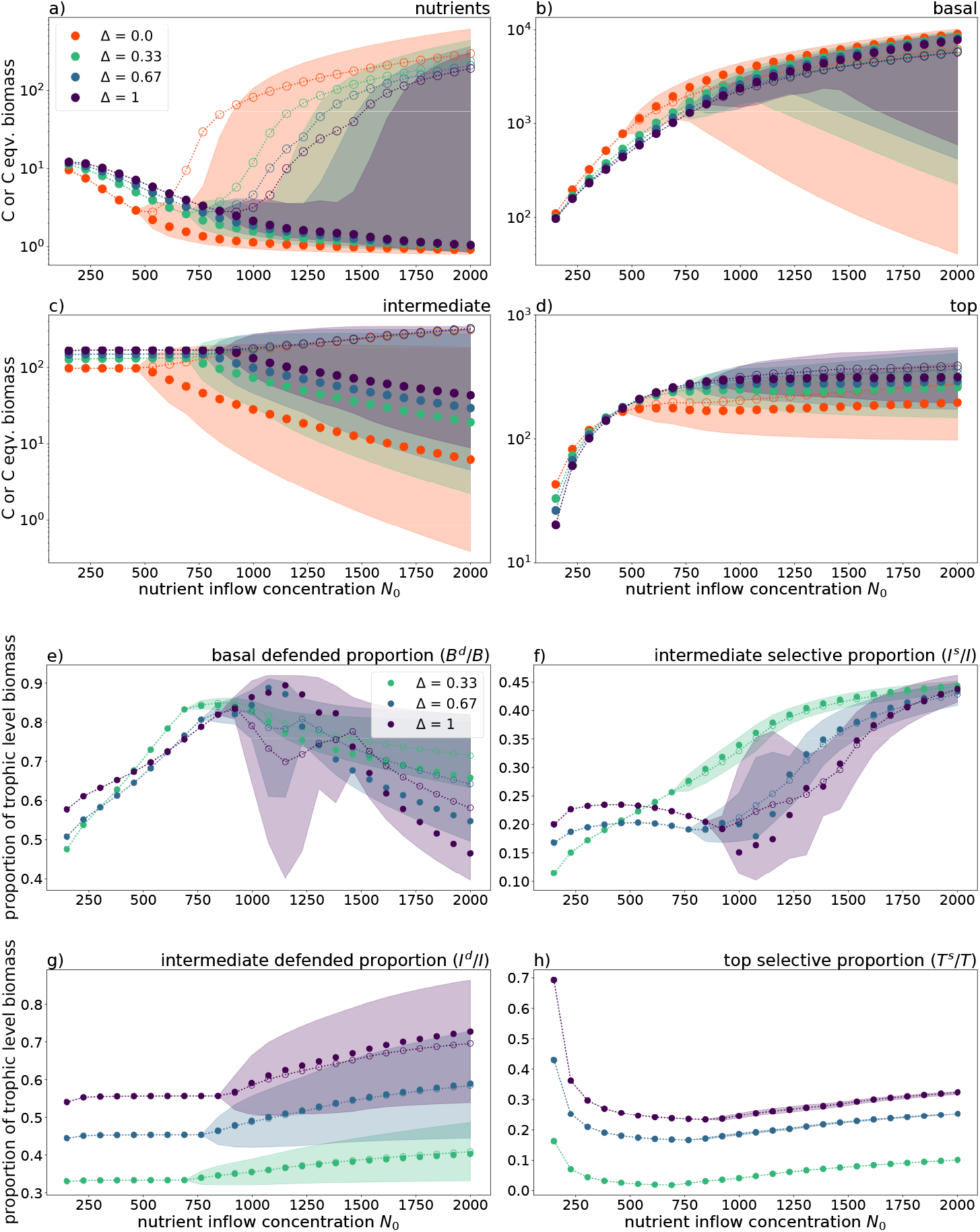
a) to d): Mean (empty dots connected by dotted line), median (solid dots), and interquartile range (shaded) of biomass dynamics across trophic levels across a range of *N*_0_; e), g): proportion of basal/intermediate trophic level exhibiting defence; f), h): proportion of intermediate/top trophic level exhibiting selectivity, over the same range of *N*_0_. The parameters used are as stated in Table 1. Please note the differences in *y*-axis scale across the panels.

**Fig. 3.**
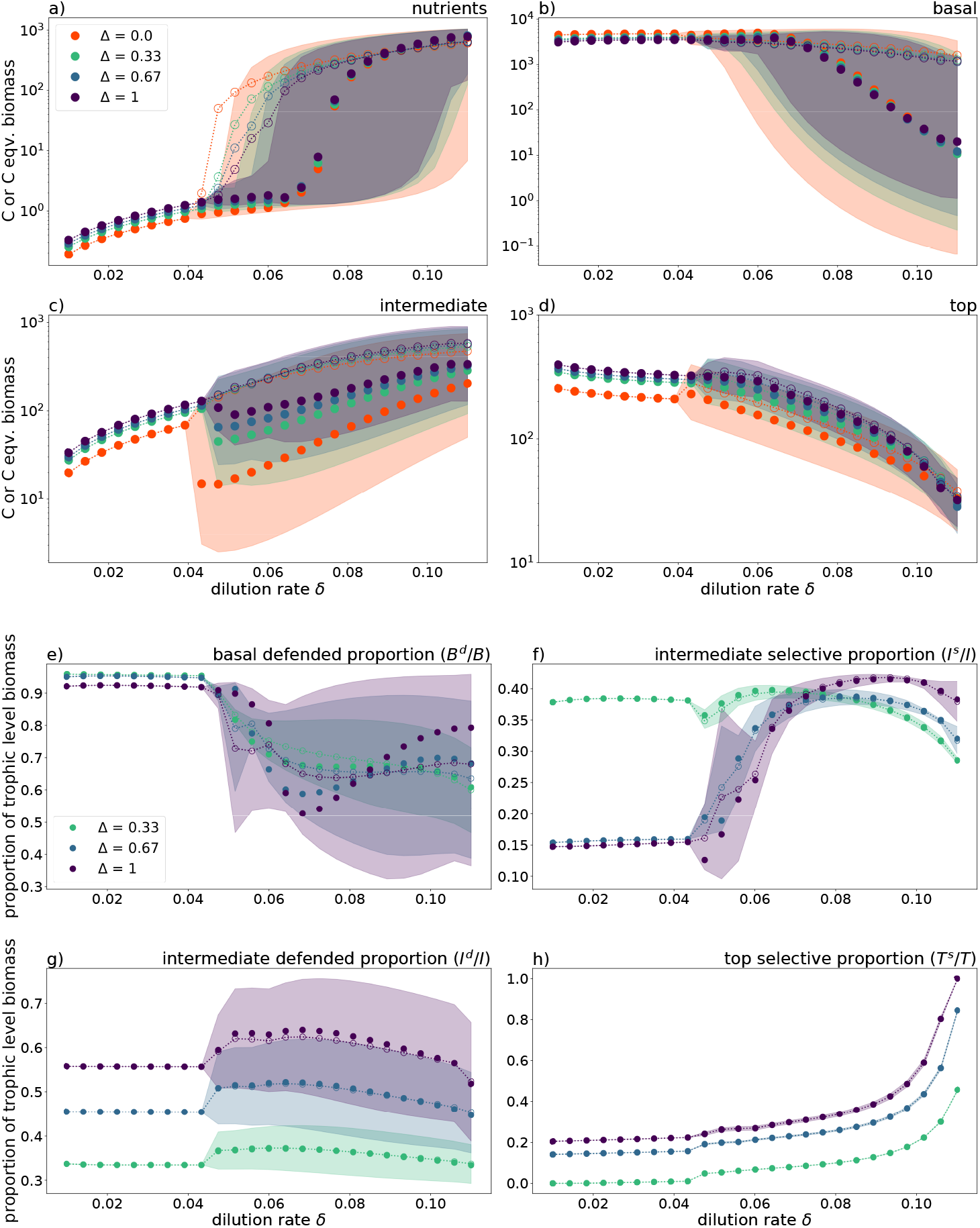
a) to d): Mean (empty dots connected by dotted line), median (solid dots), and interquartile range (shaded) of biomass dynamics across trophic levels across a large range of *δ*. e), g): proportion of basal/intermediate trophic level exhibiting defence; f), h): proportion of intermediate/top trophic level exhibiting selectivity, over the same range of *δ*. The parameters used are as stated in Table 1. Please note the differences in *y*-axis scale across the panels.

**Fig. 4.**
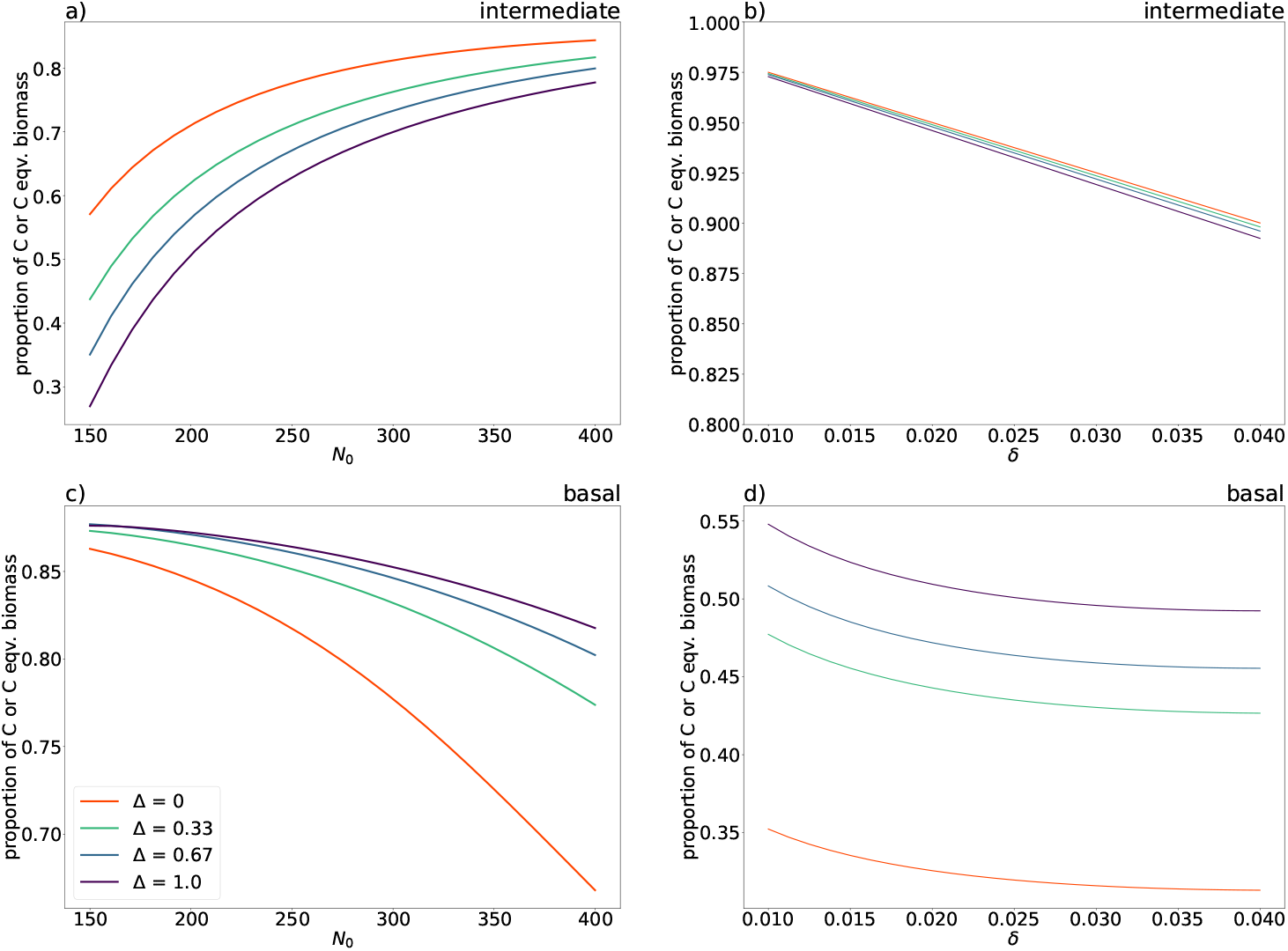
Proportion of production of one trophic level transferred up to the next trophic level for food webs of differing functional diversity, across a range of values of nutrient inflow concentration *N*_0_ (a, b) and *δ* (c, d) for which the system relaxes to a fixed point. The remaining part is washed out and replaced with nutrient-rich substrate at concentration *N*_0_. Where *N*_0_ is perturbed, *δ* = 0.055 l/day. Where *δ* is perturbed, *N*_0_ = 1200*µ*g/l.

**Fig. 5.**
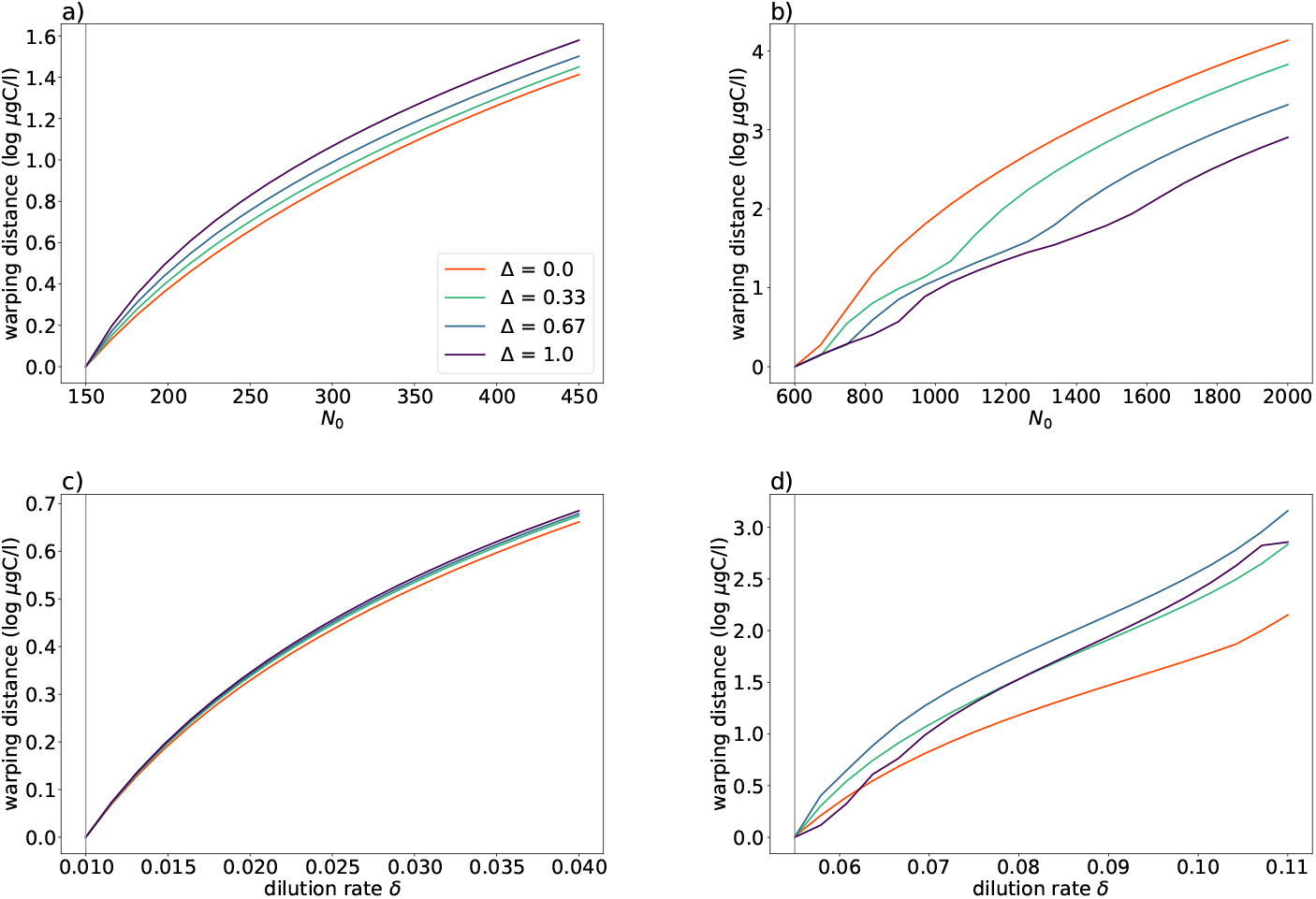
Average warping distances between pre- and post-perturbation attractors for webs of varying degrees of functional diversity. Panels (a) and (b) show the average warping distances when *N*_0_ and *δ* respectively are perturbed within the fixed point regime; panels (c) and (d) correspond to perturbations to *N*_0_ and *δ* respectively within the limit cycle regime. In each case, the pre-perturbation values of *N*_0_ and *δ* are indicated by grey lines.

**Fig. 6.**
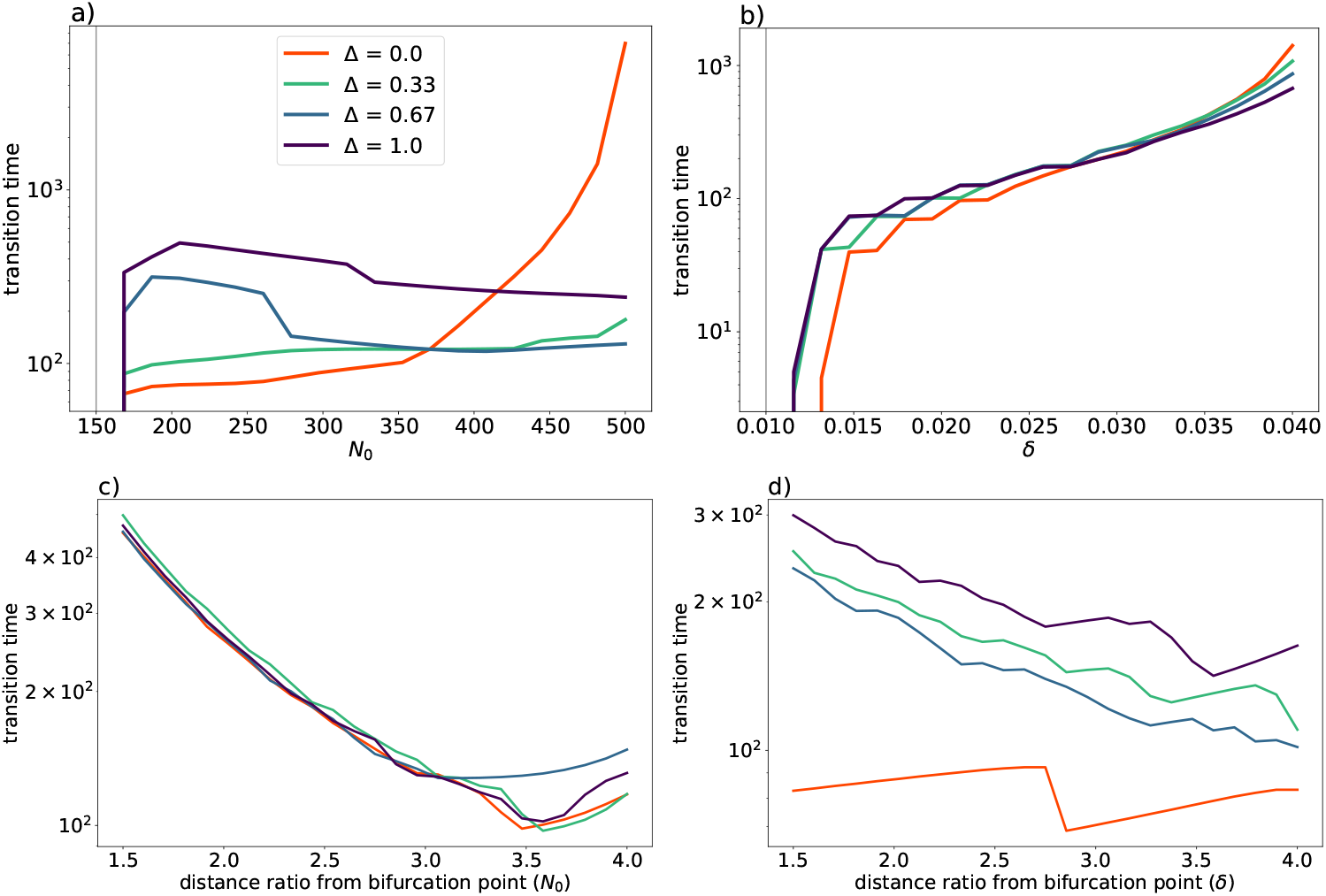
a), b): Transition times between pre- and post-perturbation fixed points for systems spanning the full range of functional diversity, for perturbations to *N*_0_ (a) and *δ* (b). The pre-perturbation value is indicated by a grey line. c), d): Transition times between pre-perturbation attractors very close to the bifurcation and post-perturbation attractors for values of *N*_0_ (c) and *δ* (d) which are 1.5 to 4 times lower than the bifurcation value.

**Fig. 7.**
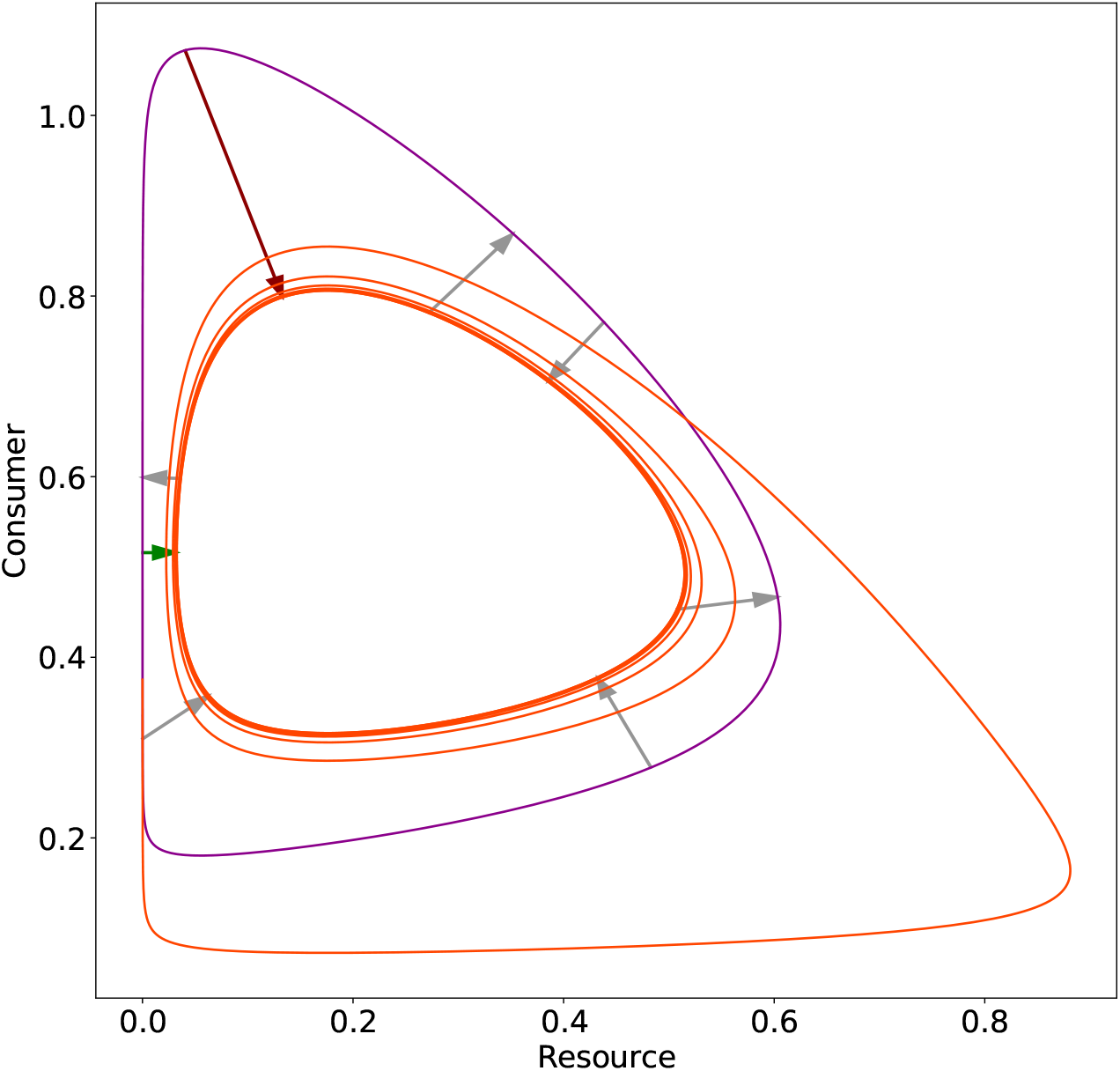
An illustration of the method of calculating the warping distance. Two Rosenzweig-MacArthur attractors are shown - the pre-perturbation attractor in violet, and the post-perturbation attractor in orange. The trajectory of the system from the pre- to the post-perturbation attractor is shown as a dotted orange line. The shortest (largest) distance from any point on one attractor to its nearest neighbour on the other is shown in green (red, this corresponds to the maximum warping distance as defined in [Raatz et al., 2019]). A random selection of six distances from other points on one or the other attractor to their nearest neighbours on the opposite attractor are given in grey. The average warping distance is the average length of all possible arrows between points and their nearest neighbours. For details, see Methods.

The average warping distance between two attractors A and B is the average distance between nearest neighbour pairs (*p*_*A*_, *p*_*B*_) and (*q*_*B*_, *q*_*A*_). The average warping distance lends itself to an interpretation as the ‘average smallest difference in biomasses between the points on attractor A and on attractor B. In contrast to [Raatz et al., 2019], the Manhattan distance or *L*_1_-norm was used as the distance metric, in order to allow a more ecologically intuitive interpretation of the measure. A justification for this change is provided in the Appendix.

As the warping distance is a measure purely of the distances between nearest neighbour points in phase space and does not consider the evolution in time of trajectories along the attractor, any differences in the time evolution of the attractors - frequency or period, most notably - are not reflected by any change to the warping distance.

#### 2.3.3 Duration of transient

A final measure of interest was that of the *transition time* of the system, which is the time required for the system to settle on its new equilibrium. Due to difficulties in numerical computation, only transition times to stable fixed points were investigated. To compute the transition time of the system to its new fixed point, the post-perturbation trajectory was examined to locate the time at which the trajectory reaches and remains within a Manhattan distance of 10 *µgC/L* of the post-perturbation fixed point. 10 *µgC/L* corresponds to approximately one per cent of the total carbon (or carbon-equivalent in the case of nutrients) mass of the system. The time taken for the trajectory to reach this point was used as a measure of the transition time.

## 3 Results

### 3.1 Biomass dynamics

We present here the responses of a tritrophic food web with variable levels of functional diversity at each trophic level as nutrient inflow concentration *N*_0_ (Fig. 2) or dilution rate *δ* (Fig. 3) were perturbed. We began by examining the system at low levels of these parameters, for which the model settled on static equilibria. We then continued to increase the perturbation parameters past the point at which the system underwent a Hopf bifurcation and into the region for which it exhibited dynamic equilibria (limit cycles). Consistent with the predictions of classical food web theory, we observed a trophic cascade. As increases in *N*_0_ and in *δ* correspond respectively to increased bottom-up and top-down forcing, these cascading effects were opposite for the two environmental changes.

An increase in *N*_0_, implying a higher supply of nutrients increased the biomass of the basal species (Fig. 2b). The intermediate species therefore grazed more and produced more - but rather than this leading to an increase in intermediate biomass, the increased intermediate production was immediately grazed down by the top species (Fig. 4a), which in turn increased its biomass under a constant mortality rate (Fig. 2d).

Note that after an increase in *N*_0_ while the system exhibited limit cycles, mean intermediate biomass slightly increased (Fig. 2c). However, over the same range of *N*_0_, median intermediate biomass decreased. The information provided by the combination of these summary statistics suggested that the biomass remained approximately constant with increasing *N*_0_, as observed for the fixed point regime, although the amplitudes of oscillation increased.

On the other hand, an increase in *δ* impacted the top species most heavily, which explains the strong decrease observed in top biomass (Fig. 3d). Thus, the intermediate species was released from predation pressure and grazed more on the basal species. This led to an increase in intermediate biomass (Fig. 3c) and a decrease in basal biomass (Fig. 3b).

The cascading effects described above predict that the top biomass increases and intermediate biomass remains constant with an increase in *N*_0_ (Fig. 2d) and top biomass decreases and intermediate biomass increases with an increase in *δ* (Fig. 3d). However, functional diversity (Δ) had direct effects on these cascades. Increasing Δ inhibited the changes in median intermediate biomass which occurred as a result of increasing *N*_0_ and *δ*, in contrast mean intermediate biomass remained largely unaffected by Δ. (Figs. 2c and 3c). Higher levels of Δ supported higher mean and median levels of biomass at the top trophic level across the majority of the ranges of *N*_0_ and *δ* considered (Figs. 2d and 3d). Consequently, functional diversity enhanced the maintenance of top trophic level biomass – thus decreasing or increasing the resistance of top trophic level against increase in *N*_0_ or *δ* respectively– and increased the resistance of the intermediate trophic level.

### 3.2 Trait dynamics

Considering the trait dynamics at each trophic level helps to understand the biomass dynamics. Low levels of *N*_0_ mean a scarcity of nutrients, and a low production at the basal level, and consequently also at the intermediate level. At *N*_0_ = 150 *µ*gN/l, low intermediate production favoured the selective top species, with its lower half-saturation constant (Fig. 2h). As *N*_0_ increased, this advantage fell away: the proportion of nonselective top species grew markedly, which was matched by a pronounced increase in the intermediate production passed to the top trophic level (Figs. 2h and 4a).

High levels of *δ*, corresponding to high levels of natural mortality, require high growth rates to compensate for high losses. Since undefended intermediate species were available across the range of *δ* considered (Fig. 3g), selective top species therefore had an advantage over non-selective top species due to a lower half-saturation constant, and therefore a more quickly maximised growth rate. These effects were more pronounced with higher Δ. That is, the more extreme selectivity possible in more functionally diverse webs (higher Δ) allowed for greater advantage under these scarce nutrient or high mortality conditions (Figs. 2h and 3h).

In cases of larger changes to *N*_0_ and *δ*, there was a trend towards a lower mean and median defence in the basal species and a corresponding increase in mean and median selectivity in the intermediate species (Fig. 2e-f for *N*_0_ *>* 700 *µ*gN/l and Figure 3e-f for 0.04 *< δ <* 0.06 l/day). In contrast to the fixed point regime, this change was more pronounced in webs with higher Δ when the system exhibited limit cycles.

### 3.3 Trophic transfer efficiency

At very low *N*_0_ values (ca. 150 *µ*gN/l) the food chain (Δ = 0) supported more top biomass than the fully diverse food web (Δ = 1), but at higher levels of *N*_0_ (ca. 400-500 *µ*gN/l) the opposite was the case. That is, there was a more efficient conversion of increases in *N*_0_ to top biomass as functional diversity increased: for the top trophic level, at all nutrient inflow concentrations, the gradients of the efficiency curves (Fig. 4c) for higher levels of Δ were larger than those for lower levels of Δ.

For webs of higher functional diversity, a larger proportion of basal biomass reached the intermediate level as *N*_0_ increased within the fixed point range (Fig. 4c). This implies that the presence of diversity allowed the intermediate species to better overcome the defence of the basal species within this range. Supporting this is the observation that higher functional diversity webs saw a slower increase in the proportion of defended basal species as *N*_0_ increased within the fixed point regime (Fig. 2e, *N*_0_ *<* 750*µgN/l*). In conclusion, the lower resistance of the top trophic level against an increase in *N*_0_ in diverse systems emerged from a more efficient biomass transfer towards higher trophic levels due to a higher ability of the intermediate species to overcome the basal defence.

### 3.4 Onset of cyclic behaviour and oscillatory amplitudes

The onset of oscillations in the system occurred at higher values of *N*_0_ and of *δ* as Δ increased, although the phenomenon was less pronounced under perturbations to *δ*. Oscillations began in the chain (Δ = 0) at just below *N*_0_ = 500*µgN/l* and at just below *δ* = 0.04*l/day*, and in the fully diverse web (Δ = 1) shortly after *N*_0_ = 800*µgN/l* and *δ* = 0.046*l/day* (Figs. 2 and 10). Therefore, if the system began at equilibrium with a value of *N*_0_ less than 400 *µgN/l* or *δ <* 0.04*l*/day, and a perturbation was applied which increased either of these parameters, a more functionally diverse system would be less likely to begin exhibiting oscillatory dynamics than a less diverse system.

**Fig. 8.**
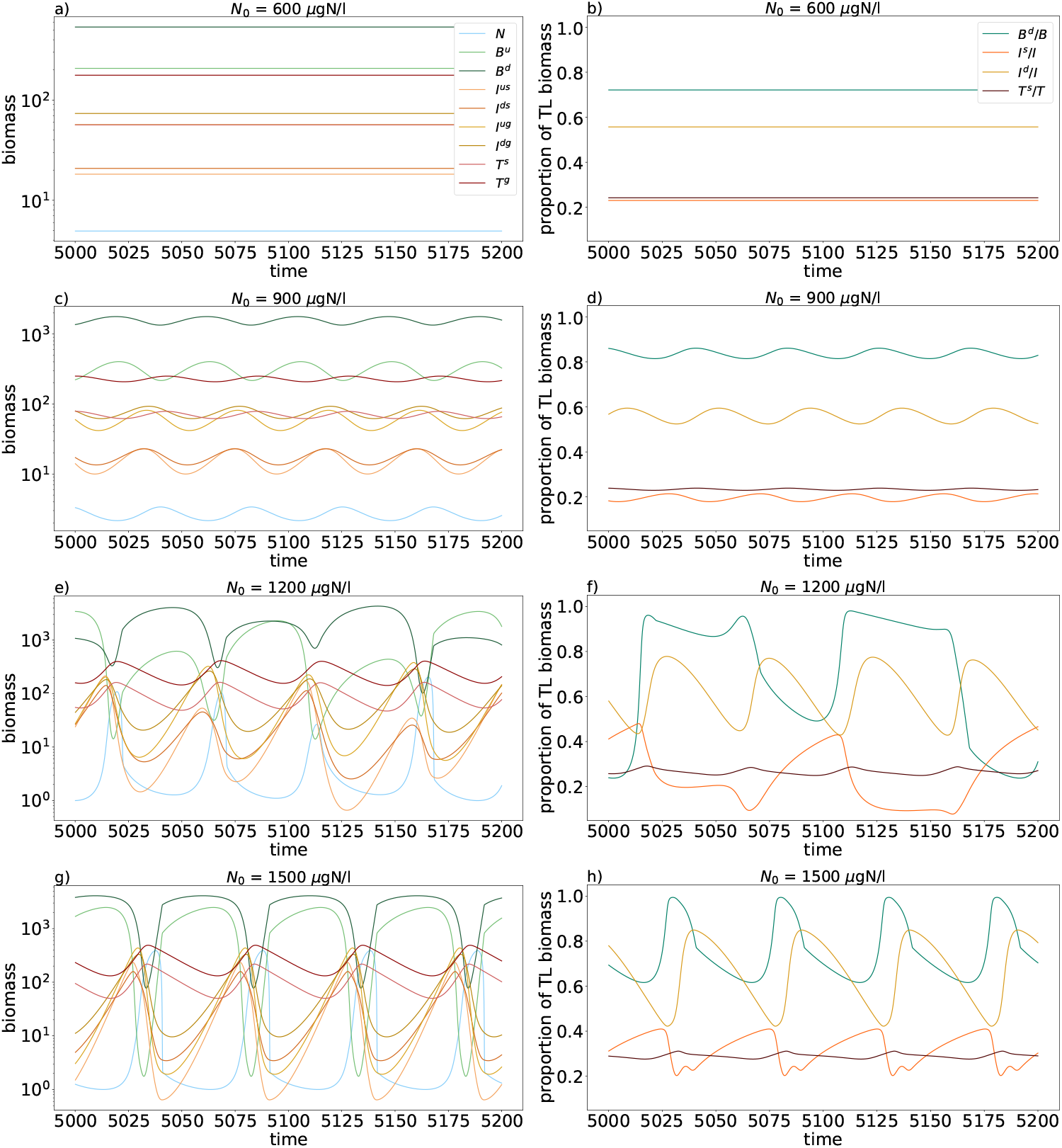
Examples of biomass (a, c, e, g) and trait (b, d, f, h) time series at static (a, b), and dynamic (c to h) equilibrium at increasing levels of *N*_0_. In particular, at panels e and g, larger amplitude oscillatory attractors exhibited long intervals at low nutrient concentrations punctuated by brief spikes at higher concentrations. This was complemented by long intervals of very high basal biomass concentrations punctuated by brief dips to markedly lower concentrations. Panels e and f illustrate the interplay between basal (B) defence and intermediate (I) selectivity at intermediate levels of *N*_0_. Throughout this figure, Δ = 1 and *δ* = 0.055 l/day. u = undefended; d = defended; s = selective; and n = nonselective, so that, for example, *T* ^*s*^*/T* denotes the proportion of the top (T) trophic level exhibiting selectivity, etc.

**Fig. 9.**
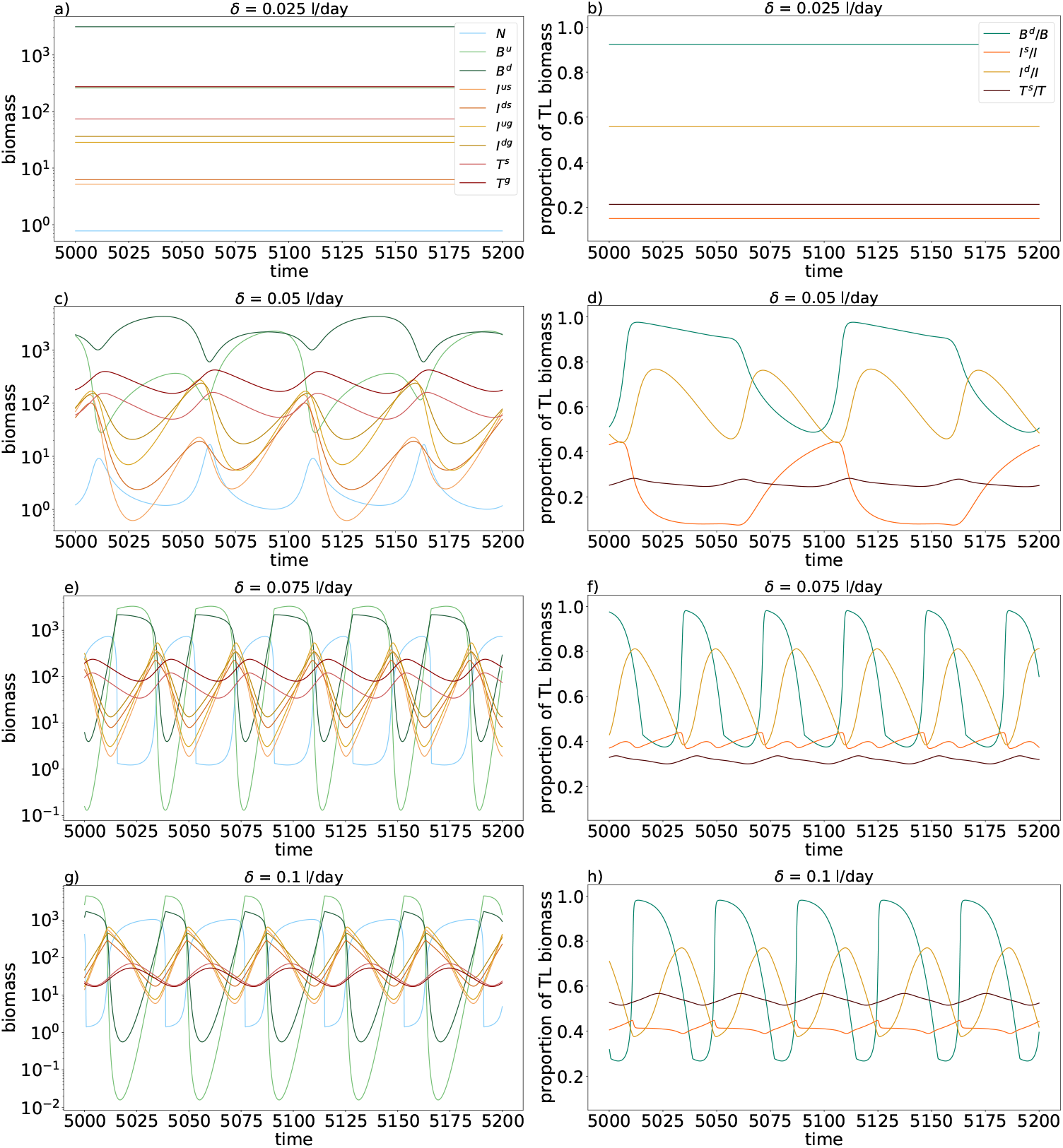
Examples of biomass (a, c, e, g) and trait (b, d, f, h) time series at static (a, b), and dynamic (c to h) equilibrium at increasing levels of *δ*. In particular, panels c and d illustrate the interplay between basal (B) defence and intermediate (I) selectivity at lower intermediate levels of *δ*, and a comparison of panels f and h illustrates the rapid rise of top selectivity as *δ* was raised to extreme values. Throughout this figure, Δ = 1 and *N*_0_ = 1200*µgN/l*. u = undefended; d = defended; s = selective; and n = nonselective, so that, for example, *T* ^*s*^*/T* denotes the proportion of the top (T) trophic level exhibiting selectivity, etc.

**Fig. 10.**
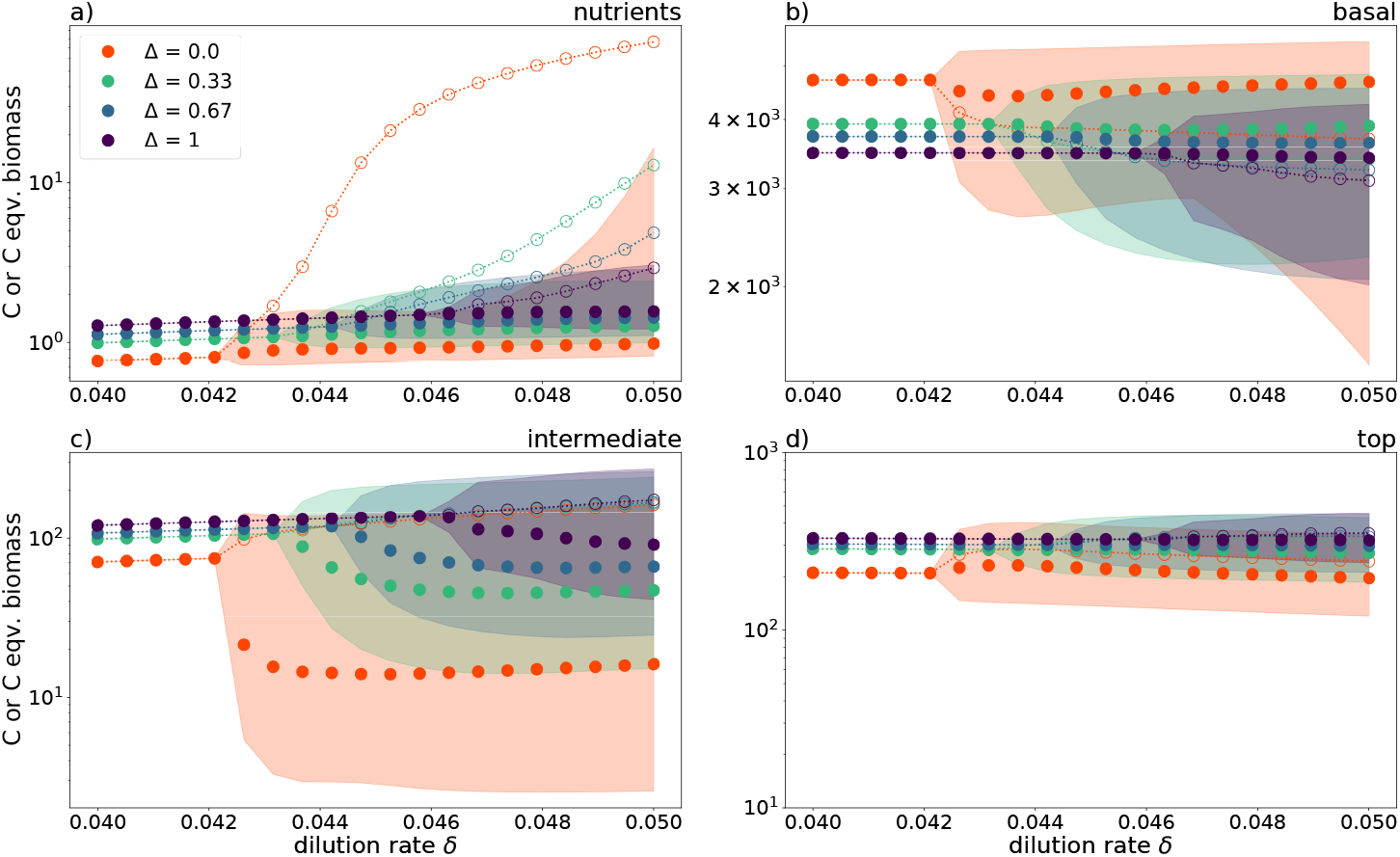
Mean (empty dots connected by dotted line), median (solid dots), and interquartile range (shaded) of biomass dynamics across trophic levels across a range of *δ* small enough for the delayed onset of oscillations due to functional diversity to be clearly visible. The parameters used are as stated in Table 1. Please note the differences in *y*-axis scale across the panels.

Moreover, should the system begin cycling as a result of the perturbation, then the higher the degree of functional diversity, the smaller would be the amplitudes of the ensuing oscillations. Indeed, the interquartile range of biomass at every trophic level was smaller for higher levels of Δ. In particular, for high *N*_0_, the chain exhibited oscillations of basal and intermediate species with an interquartile range roughly an order of magnitude larger than that of the fully diverse web (Figs. 2 and 3). This would correspond, naturally or *in vitro*, to lower functional diversity systems having a higher probability of extinctions occurring near biomass minima.

In contrast, food webs of increasing functional diversity exhibited larger amplitude temporal fluctuations in the proportion of selectivity and defence at each trophic level (Figs. 2 and 3 e-h), while exhibiting smaller amplitude temporal fluctuations in overall trophic level biomass (Figs. 2 and 3 a-d). Figures 8 and 9 in the Appendix illustrate the biomass and trait dynamics of the fully diverse web (Δ = 1) for different perturbations to *N*_0_ and *δ*.

Taken together, these observations demonstrate that the increased flexibility afforded by increased functional diversity contributed to a reduction in the overall temporal variability of a trophic level via oscillatory trait dynamics (i.e. changes in the proportion of species exhibiting a trait) within the trophic level.

### 3.5 Warping distances

The warping distance provides a summary of the overall difference between two attractors. In all cases, larger perturbations gave rise to larger warping distances, but diversity had different effects on warping distances depending on the environmental parameter and the size of the perturbation (Fig. 5).

When increasing *N*_0_ from 150 *µgN/l* a negative diversity effect was observed (Fig. 5a). Functional diversity increased the warping distances between the pre- and post-perturbation fixed points: the chain (Δ = 0) experienced less change in this region than the more diverse (Δ *>* 0) systems. A clear positive diversity effect - that is, that more diverse food webs had smaller warping distances - was observed after increasing *N*_0_ beyond 600*µgN/l* (Fig. 5b) into the limit cycle regime. Here, systems with higher levels of functional diversity tended to settle on post-perturbation states which were closer to their pre-perturbation equilibria than those of lower diversity systems.

No diversity effect was observed when perturbing *δ* from 0.015 l/day to values within the fixed point regime (Fig. 5c). Increasing *δ* from 0.055 l/day had a ‘binary’ negative effect: the chain experienced less change than the more diverse webs, all of which responded similarly (Fig. 5d). Overall, the resistance to press perturbations was context-dependent when measured by the warping distance.

### 3.6 Transition times

This subsection examines estimates of the time needed by the system to settle on its post-perturbation attractor - the *transition time* of the system for a given perturbation. This measure could be calculated only in the case of fixed point regimes.

The general observed pattern was that systems with higher values of Δ exhibited longer transition times than systems with lower Δ. This is clearly visible in Figure 6a, across all levels of functional diversity for post-perturbation values of *N*_0_ up to 350 *µgN/l*. While transition times for perturbations to *δ* were much more uniform across levels of functional diversity, the same overall pattern held: in particular, at *δ* ≤ 0.025 l/day, the most diverse system (Δ = 1) generally had slightly longer transition times than the least diverse system (Δ = 0).

As *N*_0_ increased past 350 *µgN/l* and approached 500 *µgN/l*, transition times for the chain began to increase markedly. Figure 2 illustrates the reason for this increase: as *N*_0_ approached 500 *µgN/l* from below, the stable state approached a Hopf bifurcation. In general, as a system approaches such a bifurcation point, the speed of its approach to its stable state slows exponentially. The transition times of the chain (Δ = 0) were therefore markedly higher than those of the more diverse food webs not due to Δ inherently, but because the system was experiencing so-called *critical slowing-down* as it neared its bifurcation point, which occurred for Δ *>* 0 only at higher values of *N*_0_. This phenomenon was also observed as *δ* was increased (Fig. 6b for *δ >* 0.035 l/day), although the effect is less pronounced due to Δ having less effect on the bifurcation points (Fig. 10), as compared to *N*_0_.

To overcome this confounding phenomenon, we measured the transition times of the systems from their bifurcation points to the fixed points arising from values of *N*_0_ and *δ* which were between 1.5 and 4 times smaller than the value of the perturbation parameter at the bifurcation point (Fig. 6c-d). No clear gradient across diversity was seen for perturbations to either *N*_0_ or *δ*, but across the entire range of *δ*, the chain (Δ = 0) had consistently lower transition times than the food webs (Δ *>* 0). As the attractors moved further away from the systems’ bifurcation points, the transition times of all systems decreased and became more similar. When the systems were strongly perturbed, to values of *N*_0_ and *δ* approximately 3.5 times lower than the bifurcation value, transition times generally began to increase again. In general, more diverse systems tended to have longer transition times than less diverse systems.

## 4 Discussion

We observed that increasing functional diversity increased resistance to environmental change, modelled as an increase in nutrient inflow concentration *N*_0_ or dilution rate *δ*, by dampening cascading effects and by delaying the onset of oscillations. This is in line with previous results [Folke et al., 2004, Raatz et al., 2019], although studies differ in the type of perturbation applied and in the food web structure. Functional diversity also maintained a higher top biomass, which implies a lower resistance due to a more efficient nutrient transfer to the top trophic level with increases to *N*_0_. Furthermore, following an environmental change, the warping distances between attractors were context-dependent, and higher diversity food webs generally had longer transition times than lower diversity food webs.

A study of the relationship between horizontal diversity (i.e. the diversity within each trophic level) and recovery rate in the context of herbicide or pesticide exposure showed that tritrophic food webs with higher horizontal diversity recovered more quickly after pulse perturbations [Zhao et al., 2019]. Similarly, using the same model as the present study, with a slightly different parameterisation, a higher functional diversity increased the recovery rate and resistance of food webs after a nutrient pulse [Wojcik et al., 2021].

When analysing the summarised dynamics in Figures 2 and 3, we considered both mean and median, as well as the interquartile range, with the aim of providing a comprehensive but concise summary of the dynamics: while the mean is perhaps a more obvious summary statistic, the median provides information as to the shape of the time series; it is less affected by occasional short but pronounced spikes in biomass. The nutrient level, for example, was characterised by long periods at low concentrations, interrupted by periodic spikes to values several orders of magnitudes higher. The amount of time spent at low concentrations caused the median to be comparatively low; the amplitude of the spikes was sufficiently high to cause a marked increase to the mean (Fig8e, g). This demonstrates that considering only the mean can be misleading when analysing systems in which strong oscillations occur frequently.

### 4.1 The role of the basal-intermediate interaction in resistance

We found that the basal-intermediate interaction played an important role in explaining the resistance of the food web, in line with [Wojcik et al., 2021]. In cases when the intermediate species’ ability to keep the basal species under top-down control was weak, the resistance of the food web was low. This eventually led to the extinction of the undefended basal species, and at times even to the extinction of the complete basal level, and subsequently the rest of the food web. In our study, increasing functional diversity Δ caused larger oscillations in the mean traits, in turn leading to smaller oscillations of total trophic level biomass as *N*_0_ and *δ* were perturbed.

The phenomenon was particularly strongly observed for intermediate levels of *N*_0_ and of *δ*, for which large oscillations in the proportion of defended basal species (between 40 and 90 per cent) and selective intermediate species (between 10 and 35 per cent) allowed for very small oscillations, of less than one order of magnitude, in total trophic level biomass. This pattern in the traits and biomasses remained observable as *N*_0_ and *δ* were individually increased to more extreme values. This means that adaptation through a change in the proportions of these traits could buffer the effects of eutrophication (in the case of perturbations to *N*_0_) and of increased natural mortality, brought about, for example, by increasing temperature (in the case of *δ*) by reducing the oscillations of total trophic level biomass. This suggests a sensitive coupling between the traits: small changes in biomass drove large trait changes, which in turn acted as a negative feedback mechanism for the changes in biomass - contributing, overall, to the system’s resistance to environmental change.

At much higher *N*_0_, less diverse food webs had a higher proportion of defended basal species for approximately the same proportion of selective intermediate species (Fig. 2 e, f). Additionally, less diverse food webs supported slightly higher levels of basal biomass. In other words, in less diverse food webs, a smaller proportion of basal biomass was under top-down control and available for intermediate level grazing. Such considerations illustrate the need to consider multi-trophic webs: the degree of basal top-down control depends on the level of intermediate biomass, which in turn is influenced by the top trophic level.

### 4.2 Eutrophication and energy flux

Eutrophication, i.e. increased nutrient influx and concentrations, is known to promote the development of harmful algal blooms, which can be due to an excess of inedible, defended algae [Heisler et al., 2008, Sommer et al., 2012, Selander et al., 2019]. As our systems became more eutrophic, less diverse food webs exhibited more rapid increases in the proportion of defended basal species, and also larger amplitudes of oscillations in overall basal biomass (Fig. 2b). That is, less diverse herbivores exerted less top-down control. This suggests that in less diverse food webs, harmful algal blooms may occur for longer, and with higher magnitude. This study thus highlights that integrating functional diversity to cope with eutrophication may improve management strategies.

Besides the occurrence of harmful algal blooms, other consequences of eutrophication in theoretical models are an increase in top biomass and larger amplitude oscillations in biomasses at the species level [Azzali et al., 2017]. This increase in oscillatory amplitude is expected by the paradox of enrichment [Rosenzweig, 1971] when trait dynamics are not taken into account. However, trait variation played an important role in stabilising the dynamics of bitrophic systems [Vos et al., 2004, Mougi and Nishimura, 2008], including in more complex models with spatial resolution [Azzali et al., 2017]. Our study supports these results, and emphasises that higher functional diversity lessens the destabilising effects of eutrophication on the dynamics of a tritrophic food web (Fig. 2). On top of an increase in amplitudes of oscillation of species biomass, eutrophication led, as expected, to an increase in top biomass, in line with other theoretical and empirical studies [Hulot et al., 2000, Wollrab et al., 2012].

Our results correspond partly to the principle of energy flux developed by [Rip and McCann, 2011], which predicts that an increase in energy flux to consumers will destabilise their dynamics, insofar as an increase in *N*_0_ was observed to increase the amplitudes of oscillations. We found that functional diversity increased the efficiency of the systems, in effect increasing the top biomass which is expected to destabilise dynamics [Rip and McCann, 2011]. However, the destabilising effect of eutrophication (i.e. an increase in the amplitude of biomass oscillations) was more effectively dampened via trait dynamics when functional diversity was higher. In turn, functional diversity had a general stabilizing effect (i.e enhanced resistance) since the stabilizing effect via the dampening of the biomass oscillations overruled the destabilizing effect due to the increased biomass transfer to the top. Accounting for functional diversity illustrates the boundaries of the applicability of the principle of energy flux in its present form.

### 4.3 Temperature change

In this study, perturbations to dilution rate *δ* were used to model environmental change which affects the top level most severely. Experimental and theoretical work has shown that increased temperature primarily impacts the top of planktonic food webs, and that intermediate trophic levels are more directly impacted by altered trophic interactions than by an increase in temperature [Binzer et al., 2012, Binzer et al., 2016, Murphy et al., 2020]. Perturbing *δ* therefore allowed us to investigate possible effects of warming without having explicit control over temperature in our model.

Warming has been shown to increase the metabolic rate of predators, but may not correspondingly increase their rate of ingestion [Rall et al., 2010]. Therefore, in order to survive warming, top predators must adapt in some other way to overcome this imbalance. The current study suggests a possible mechanism for survival in such a situation, dependent on the presence of functional diversity. The top predators can maximise their grazing rate, and therefore their growth rate, by lowering their half-saturation constant. This comes at a cost of limiting its range of prey via increased selectivity. We observed that, as *δ* increased to extreme values, the top trophic level became steadily dominated by selective predators - up to 100% when Δ = 1 for the highest dilution rates investigated - enabling the top level to maximise its growth rate (Fig. 3h). This led to maximum growth rates at the top level which could compensate for the high mortality rate imposed on the system by increased *δ*. For higher-diversity systems, this occurred despite the dominance of defence at the intermediate level, and furthermore such systems supported a higher level of top and intermediate biomass (Fig. 3c, d). In other words, functional diversity provided a mechanism for the top trophic level to mitigate, to some extent, the negative effects of adverse environmental conditions, such as could be brought about by warming.

### 4.4 Warping distances

The warping distance was first used in an ecological context in [Raatz et al., 2019] to measure the overall change in an attractor caused by a press perturbation: a bitrophic system’s resistance to perturbations to nutrient inflow concentration increased with trait adaptation speed. The current investigation builds on this result in the following ways. Firstly, we extended the application of the measure from a bitrophic to a tritrophic model, and confirmed the previous finding in the case of eutrophication. Furthermore, the present study argues that, in an ecological context, the Manhattan distance is a preferable metric to the previously used Euclidean distance for measuring distances between points in biomass phase space. A full argument is provided in section A1 of the Appendix.

Species richness increases the likelihood of a system displaying a variety of diversity-stability relationships [Ives and Carpenter, 2007]. In line with this, we found highly context-dependent variation in the relationship between functional diversity and warping distance post-perturbation. Other measures studied in this investigation allowed for a more granular analysis of the dynamics; we suggest that the warping distance is a too aggregated measure to deliver mechanistic insights. The trophic level with the largest scale factor change in biomass dominates the measure - in our case, the basal species dominates the warping distance.

### 4.5 Transition time

The transition time measures the time needed by the system to settle on its new attractor after a press perturbation. It is similar to, but differs slightly from other temporal measures found in the literature such as the return time or the recovery time [Ives and Carpenter, 2007, Oliver et al., 2015]. Return and recovery times measure the elasticity of the system, i.e. the speed with which a system returns to its pre-perturbation state after a pulse perturbation. However, press perturbations cause the system’s attractor to change - and the transition time quantifies the duration of the transient as the system moves from the original to the altered attractor.

The current investigation supports previous findings that transients are highly dependent on other factors not directly under scrutiny here [Wojcik et al., 2021]. Notably, the proximity of the perturbation parameter to a bifurcation point had a markedly stronger effect on the transition time than functional diversity, and partly masked the effect of the size of the perturbation. This is in line with previous studies discussing long transients in the vicinity of bifurcation points [Hastings et al., 2018].

When systems of different degrees of functional diversity were at the same relative distance from a bifurcation point, increased functional diversity tended to increase transition times, analogous to theoretical work on the effect of species biodiversity and return times [Ives and Carpenter, 2007]. When viewed in conjunction with warping distances, despite context-dependent variation in the relationship between functional diversity and warping distance, the change of attractor tended to occur at a gentler speed in more functionally diverse systems. This phenomenon may be because as functional diversity increased, the minimum growth rates of the species decreased. In other words, the slowest-growing species in higher-diversity webs grew more slowly than the slowest-growing species in lower-diversity webs, thus increasing the time taken for higher-diversity webs to reach their attractors.

## 5 Conclusion

Broadly, we have shown that functional diversity increased the resistance of a tritrophic food web to environmental change, modelled as press perturbations simulating eutrophication (higher *N*_0_) and increases in temperature (higher *δ*). As *N*_0_ and *δ* were perturbed, functional diversity delayed the onset of oscillatory dynamics, and buffered the amplitude of oscillations once a dynamic equilibrium had been reached due to oscillatory trait dynamics within each trophic level. We found that the effect of diversity on resistance as measured by the warping distance was context-dependent. While the lengths of the systems’ transients tended to increase with functional diversity, this diversity generally buffered the effects of press perturbations in tritrophic food webs. This highlights that in a constantly changing world, functional diversity should be considered in attempts to limit the impact of environmental stressors.

## 8 Acknowledgments

The authors would like to thank Christian Guill for stimulating comments and suggestions on the manuscript, and Ruben Ceulemans, Ellen van Velzen, and Toni Klauschies for valuable conversations and advice during each stage of the investigation. Funding was provided by the DFG (GA401/26-2).

## 9 Statements and Declarations

### 9.3 Authors’ contributions

The initial outline of the study was developed by UG. Improvement of methodology, numerical simulations and data analysis were carried out by GA. Writing lead of the manuscript and interpretation were done by GA. Further writing, interpretation, comments, and edits were done by LW. Revision of the manuscript and close project supervision was done by UG.

### 9.4 Code availability

We intend to make the code required to produce our results openly available in a public repository (Zenodo, Dryad) that issues DOIs for datasets before acceptance of our manuscript.

## 10 Appendix

### 10.1 Warping distance

The following paragraphs contain a brief justification for the use of the Manhattan distance rather than the Euclidean distance in the calculation of the warping distances between attractors in an ecological context. Figure 7 illustrates how the warping distance is computed.

The Euclidean distance between two points **u** and **v** ∈ ℝ^*n*^ is 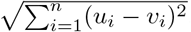, and is the usual metric by which distances are measured. While the Euclidean distance is intuitive from a purely geometric perspective as the length of the shortest straight line between two points, it is very difficult to interpret the ecological meaning of such a straight line between two points in biomass phase space. To provide a concrete example - if we are in (Resource, Consumer)-space, the Euclidean distance between A = (1, 1) and B = (2, 2) is 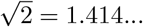, which has no clear ecological interpretation.

In contrast, the Manhattan distance between two points **u** and **v** ∈ ℝ^*n*^ is 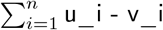, i.e. the sum of the absolute differences in position of each coordinate of the points. When working in biomass (or log biomass) phase space, the Manhattan distance can be readily interpreted as the total difference in (log) biomass over all species or over all trophic levels. Using the same example, the Manhattan distance between A = (1, 1) and B = (2, 2) is 2, which reflects the fact that to get from A to B it is necessary to grow 1 unit of resource biomass and an additional 1 unit of consumer biomass.

### 10.2 Time series of biomass and trait dynamics

This subsection is a short description of the four main types of equilibrium dynamics observed in the model with the parameterisation used in this investigation when Δ = 1. With *δ* at an intermediate value of 0.055 l/day, low levels of *N*_0_ (600 *µgN/l*) caused the system to settle on a fixed point, for which the basal trophic level was predominantly defended, and the intermediate and top levels both predominantly nonselective. Intermediate defence was relatively balanced (Fig. 8a-b). A very similar result was observed with *N*_0_ at an intermediate value of 1200*µgN/l*, low levels of *δ* (0.025 l/day) (Fig. 9a-b). Raising *N*_0_ just past its bifurcation point (and keeping *δ* constant) caused low-amplitude oscillations with similar trait ratios, notwithstanding a slightly higher proportion of basal defence (Fig. 8c-d). In contrast, larger amplitude oscillations were quickly brought on when *δ* was raised past its bifurcation point. Here the system oscillated with periods comprising two biomass peaks, one for which the basal species was predominantly defended and the intermediate species predominantly nonselective, the other of which exhibited a greater balance between these traits (Fig. 9c-d).

Further increasing *N*_0_ (1200 *µgN/l*) with constant *δ* brought the system into a region of large-amplitude oscillatory trait dynamics, in which basal defence was predominant for two of the four peaks comprising the total period, undefended basal species being predominant for one, and a roughly equal distribution for the remaining peak. The intermediate level was correspondingly affected, exhibiting peaks of high nonselectivity following intervals of high basal defence. The top trophic level remained predominantly nonselective and did not exhibit pronounced oscillatory trait dynamics (Fig. 8e-f). Raising *δ* further to 0.075 l/day, with *N*_0_ maintained at 1200 *µgN/l*, reduced the period of the attractor in biomass phase space and markedly reduced the amplitude of oscillations of intermediate selectivity (Fig. 9e-f).

At high levels of *N*_0_ (∼1500*µgN/l*), basal defence/intermediate selectivity trait dynamics became less pronounced and the system exhibited more regular oscillations, of a period roughly one-quarter the length of those in the highly oscillatory region for *N*_0_ described above (Fig. 8g-h). Extreme values of *δ* (∼0.1 l/day) caused an increase in the oscillations in the proportion of basal defence and a pronounced increase in the proportion of top selectivity, as well as lowering the minimum biomass concentration reached by the undefended basal species, as compared to the attractor at *δ* = 0.075 l/day (Fig. 9g-h).

